# Significant roles in RNA-binding for the amino-terminal domains of Drosophila Pumilio and Nanos

**DOI:** 10.1101/2023.10.24.563753

**Authors:** Tammy H. Wharton, Mohammad Marhabaie, Robin P. Wharton

## Abstract

The Drosophila Pumilio (Pum) and Nanos (Nos) RNA-binding proteins govern abdominal segmentation in the early embryo, as well as a variety of other events during development. They bind together to a compound Nanos Response Element (NRE) present in thousands of maternal mRNAs in the ovary and embryo, including *hunchback* (*hb*) mRNA, thereby regulating poly-adenylation, translation, and stability. Many studies support a model in which mRNA recognition and effector recruitment are achieved by distinct regions of each protein. The well-ordered Pum and Nos RNA-binding domains (RBDs) are sufficient to specifically recognize NREs; the relatively larger low-complexity N-terminal domains (NTDs) of each protein have been thought to act by recruiting mRNA regulators. Here we use yeast interaction assays to show that the NTDs also play a significant role in recognition of the NRE, acting via two mechanisms. First, the Pum and Nos NTDs interact in trans to promote assembly of the Pum/Nos/NRE ternary complex. Second, the Pum NTD acts via an unknown mechanism in cis, modifying base recognition by its RBD. These activities of the Pum NTD are important for its regulation of maternal *hb* mRNA in vivo.

## Introduction

Translational regulation by Pum and Nos plays an important role in Drosophila, particularly at three stages in development of the ovarian germline and the early embryo. First, the initial step in formation of an egg chamber involves the division of a germline stem cell; maintenance of stem cell status requires Pum and Nos activity (1,2). Second, from late in oogenesis to the onset of zygotic transcription in the embryo, Pum and Nos jointly regulate thousands of mRNAs to sculpt the maternal transcriptome (3). Third, Pum and Nos jointly regulate maternal *hb* mRNA in the posterior of the syncitial cleavage stage embryo to allow abdominal segmentation (4,5). In addition to these three functions, Pum and Nos govern many other biological processes in Drosophila, including various aspects of the cell biology of primordial germ cells (PGCs) (6–8), dendritic arborization of larval sensory neurons (9,10), and remodeling of the larval neuromuscular junction (11).

Although Nos and Pum may regulate some mRNAs independently of each other (3,12), in most cases they act together, binding jointly to a compound binding site in target mRNAs. This compound site, a Nanos Response Element (NRE), is comprised of adjacent Nos and Pum binding sites (NBS and PBS, respectively; see Fig. 1A) (4,5,13). The Pum RBD, which resides in the C-terminal portion of the protein, consists of 8 homologous repeats that collectively constitute a Puf domain. Pum binds on its own with high specificity and affinity to the PBS (14,15), with each of the Puf repeats recognizing one of the 8 nucleotides in the PBS (16). Unlike Pum, the Nos RBD on its own does not bind to the NBS (4,5,17). But together, the two RBDs bind cooperatively to the NRE, with Nos contributing modest sequence specificity via contacts to the degenerate NBS as well as binding energy that stabilizes the ternary Nos/Pum/NRE complex. As a result of protein-protein interactions between the Nos and Pum RBDs, the specificity of Pum for nucleotides at the 3’-end of the PBS (positions 6-8) is relaxed in the ternary complex, and this is thought to allow regulation of an expanded repertoire of mRNAs (5).

**Figure 1.**
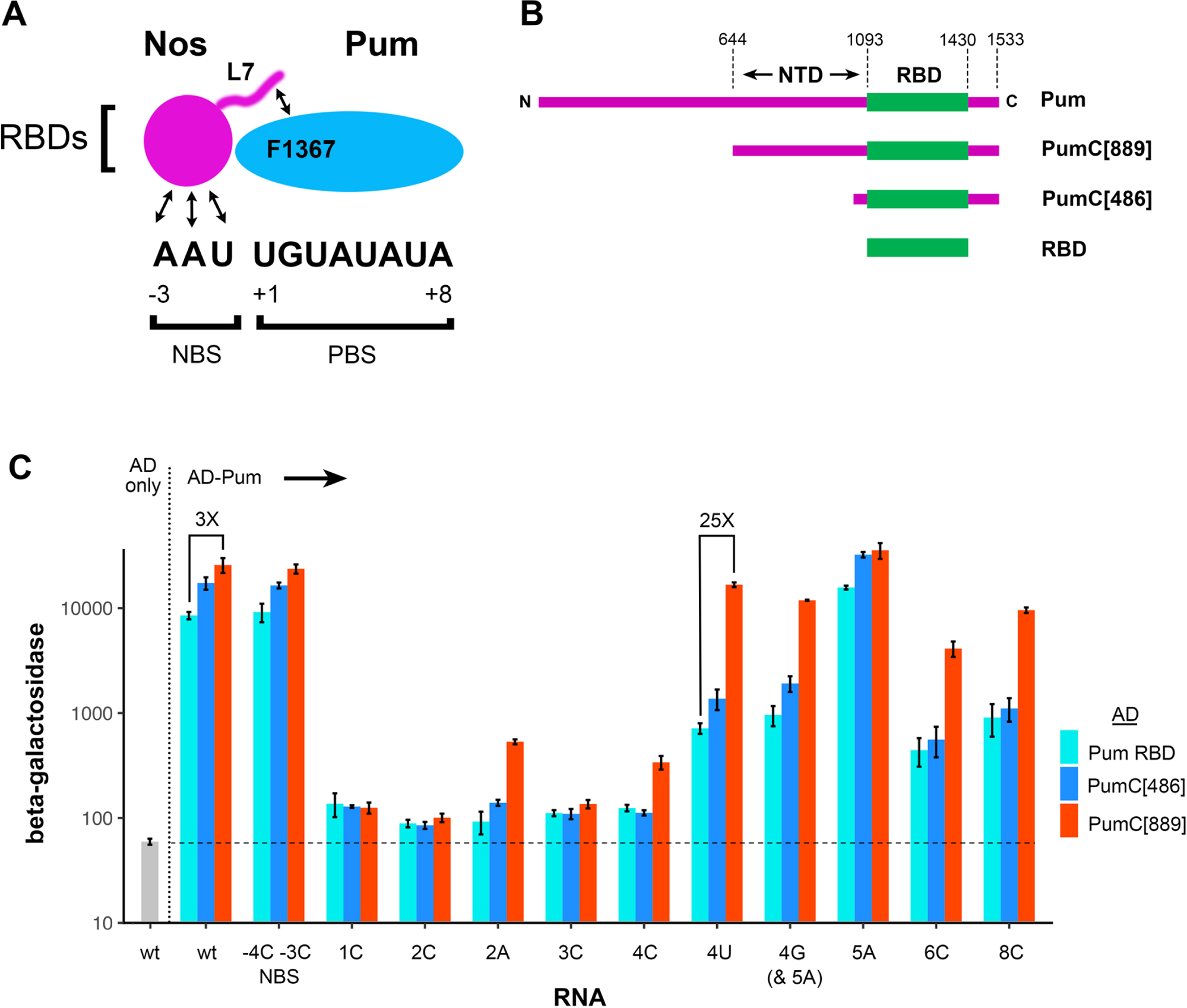
The Pum NTD alters RNA-binding activity. A. Schematic drawing of the Nos and Pum RBDs bound to the wild type NRE used in this and subsequent figures. The drawing is based on the structure of Weidmann et al. (2016), and emphasizes interactions that stabilize the ternary complex between the C-terminal tail of Nos (which is altered in the L7 mutant) and F1367 in Pum, as well as interactions of Nos with the RNA. Interactions between Pum and the RNA are omitted for clarity. B. A drawing (to scale) of proteins tested in this work, with amino acid residue landmarks above and names to the right. A major focus of this work, the Pum NTD, is a portion of the full length pA isoform of Pum, as indicated. C. Logarithmic plot of the results of β-galactosidase assays measuring binding of each of the AD-fusion proteins indicated at the right to various derivatives of a *hb* NRE in yeast three-hybrid experiments. Selected comparisons between β-galactosidase levels (based on the ratio of values in raw data) are highlighted with brackets, with the fold difference indicated above each bracket (i.e., “3X”). The y-axis is luminometer arbitrary light units (X 1000) here and in subsequent figures. Each measurement is the average of 3 or 4 independent cultures and the error bars show standard deviation. The x-axis displays the RNA site in each group of experiments here and in subsequent figures Sequence of the wild type (wt) site is in Figs. 1A and 3A, the NBS mutant has A to C substitutions at positions −4 and −3 and has been assayed for activity in vivo (14), and the remaining mutant sites are identified by position within the NRE and identity of the single nucleotide change (e.g., 1C = U to C substitution at position +1). Note that the 4G site also bears a 5A substitution to avoid creation of a fortuitous UGU sequence that would constitute the core of a cryptic Pum binding site. The empty vector “AD only” control co-expressed with the wt RNA is at the left. Although not shown, we have measured the background level of β-galactosidase in yeast co-expressing AD only with each of the RNAs; in no case is the background level of of β-galactosidase substantially different than it is with the wt RNA. The horizontal dashed line indicates the basal level of β-galactosidase here and in subsequent figures.

Both Pum and Nos are bipartite proteins, with carboxy-proximal RBDs fused to larger N-terminal domains (NTDs) that are involved in the recruitment of effectors that regulate translation and stability of bound mRNAs. The major effector recruited by Nos+Pum is thought to be the CCR4/Not/deadenylase complex (18–20). The 1091 residue Pum NTD contains three regions that can regulate bound mRNAs when tethered to a suitably engineered reporter in transfected cells; one of these regions interacts directly with the NOT1 and NOT3 subunits of the CCR4 deadenylase complex (18). The NTD of Nos also interacts with the NOT complex (20). Thus, acting via the CCR4 complex, Nos and Pum promote the deadenylation, translational repression, and ultimate degradation of target mRNAs in the embryo and in cultured cells. Both the Pum and Nos NTDs consist primarily of low-complexity sequence predicted to be intrinsically disordered, which has somewhat hindered molecular studies of their function. Although the Pum RBD may help recruit the CCR4 deadenylase complex (21), the general picture to emerge from the work summarized above is that joint mRNA target recognition by the Pum and Nos RBDs allows recruitment of effectors via their NTDs.

The extensive genetic, molecular, and structural experiments that have led to our current understanding of RNA sequence recognition by Nos and Pum has focused on studies of their structured, well-ordered RBDs. To our knowledge, no role in RNA recognition has been ascribed to the Nos and Pum NTDs.

In this report, we use yeast interaction experiments to investigate function of the Nos and Pum NTDs in RNA-binding. We find that, in addition to their previously described roles in effector recruitment, the NTDs have two important functions in target recognition. First, intermolecular interactions between the Nos and Pum NTDs play a role in RNA-binding and site selection. Second, we find a surprising intramolecular effect of the Pum NTD on the apparent affinity and binding specificity of its associated RBD. We show that this intramolecular activity of the NTD plays a role in regulation of *hb* mRNA in the Drosophila embryo.

## Results

### The Pum NTD alters RNA-binding

As described above, Nos and Pum bind jointly to a compound 11 nt RNA sequence consisting of adjacent binding sites for each protein (Fig. 1A). Formation of the ternary complex is dependent on (1) interactions between the two RBDs, involving F1367 in Pum and residues in the C-terminal tail of Nos that are deleted in the L7 mutant; and (2) protein-RNA base interactions between Nos and the NBS (4,5,22). Based on studies with the isolated RBDs, the identities of RNA bases at positions +1 through +4, UGUA, have been thought to be rigidly specified, such that substitutions at these nucleotides essentially eliminate binding of either Pum alone or joint binding of Nos+Pum.

We were therefore surprised to find that Nos+Pum jointly target mRNAs in the ovary and embryo via RNA sites bearing UGUU bases at positions +1 to +4, in addition to sites with the canonical UGUA sequence (3). In these experiments, mRNA regulation is driven by the full-length proteins Thus, one possible explanation for the apparent discrepancy between the two sets of experiments is that the Nos and Pum NTDs alter RNA site selection.

To test this idea, we first asked whether the Pum NTD has any effect on RNA-binding in yeast three-hybrid experiments (23). These measure the binding of Pum fusions with a transcriptional activation domain (AD) to various RNAs; RNA-binding of the fusion protein drives expression of a β-galactosidase reporter. In the predominant Pum isoform in early embryos, the RBD is embedded near the C-terminus of the 1533 residue protein (Fig. 1B). Pum derivatives bearing the 643 most N-terminal residues were toxic or semi-toxic in yeast; but we found that expression of a truncated derivative lacking these residues, PumC[889], had little effect on yeast viability or growth. Throughout the remainder of this report, we refer to Pum residues 644-1092 present in PumC[889] as the Pum NTD. Initially, we compared binding of PumC[889] with binding of two derivatives lacking the NTD: PumC[486] [similar to derivatives that previously have been shown to bind specifically in yeast three-hybrid experiments and to partially regulate *hb* in embryos (4)], and the minimal RBD (Fig. 1B). Expression of each protein was measured in Western blots, detecting a common epitope-tag inserted between the AD and Pum moeities. The RBD and PumC[889] are expressed at essentially identical levels, whereas PumC[486] is expressed at a 2.9-fold higher level (S1 Fig). Each Pum derivative was co-expressed in yeast with either the wild type NRE shown in Fig 1A or one of a panel of mutant sites bearing nucleotide substitutions across the 11 nt site, and β-galactosidase activity was measured to assess RNA-binding.

In previous experiments that assayed binding in vitro (5) or in the fly ovary (24), the Pum RBD has been shown to recognize the sequence UGUAHAUA (H = A, U, or G). Although we have not tested every possible substitution at each position of the NRE, our measurements of RBD binding in yeast agree with prior results. Substitutions in the core UGU sequence from +1 to +3 abolish binding, whereas substitutions at more 3’-proximal positions are tolerated (to varying extents) such that specificity at positions +4 to +8 of the NRE is partially relaxed (Fig. 1C).

In the presence of the NTD, RNA-binding is altered in two important respects. First, binding specificity of PumC[889] is further relaxed such that mutations at positions +4, +6, and +8 are well tolerated (Fig. 1C). In particular, PumC[889] binds to the 4U site only 1.5-fold less than to the wild type NRE (e.g., 4A), consistent with the observation that both canonical UGUA and non-canonical UGUU sites mediate Nos+Pum activity in vivo (3). Second, the apparent affinity of PumC[889] for the wild type NRE is elevated approximately threefold (Fig. 1C). The net result of these two alterations is substantial. For example, PumC[889] binds 25-fold better to the 4U site than the isolated RBD (Fig 1C). Finally, we note that in comparison with the isolated RBD, PumC[486] appears to have a slightly elevated affinity for the wild type NRE, but an essentially identical specificity for all mutant sites tested.

In the discussion above, we assume that β-galactosidase levels simply report RNA-binding and recruitment of the vector-encoded GAL4 AD by each Pum derivative. An alternative explanation is that the NTD bears a fortuitous transcriptional activation signal such that equivalent binding of PumC[889] and the RBD (for example) would result in higher β-galactosidase levels in the former case. As shown in S1 Data, we ruled out this possibility for PumC[889] (as well as all the other proteins tested in this report), and thus conclude that β-galactosidase levels faithfully report site occupancy and not differential activation of the LacZ reporter.

### The Pum and Nos NTDs potentiate Nos recruitment to NREs

We next asked whether the Nos and Pum NTDs affect formation of a ternary complex with NREs. To address this question, we measured Nos recruitment in yeast 4-hybrid experiments, co-expressing in the yeast strain used above various: (1) NREs, (2) Pum derivatives fused to a nuclear localization signal (NLS) but not an AD, and (3) AD-fusions to Nos (S2 Fig). Recruitment of Nos into a ternary complex was then assayed by measuring β-galactosidase levels. We once again used PumC[889] as a proxy for wild type Pum.

The Pum RBD recruits Nos into a ternary complex with the wild type NRE (Fig. 2A). Binding of Nos depends on the integrity of interactions at the RBD/RBD/NRE interface: mutations in Nos (L7), Pum (F1367S), or the NBS each independently abolish recruitment by the isolated Pum RBD, as described previously (4,5,22). However, recruitment of Nos by PumC[889] is substantially different in two respects. First, in the presence of both NTDs, recruitment of Nos to the wild type NRE is enhanced 6- to 9-fold. Second, the effect of substitutions at the RBD/RBD/NRE interface is muted; substitutions in one component of the interface (i.e., Nos or Pum or the RNA) have only a modest effect on Nos recruitment by PumC[889]. For example, the F1367S substitution reduces recruitment by PumC[889] and the Pum RBD 1.5- and 36-fold, respectively. The enhanced binding promoted by the NTDs is still sequence-specific: substitutions in both the Nos C-terminal tail (L7) and the NBS in the RNA reduce binding ∼ 100-fold, almost to background levels. Finally, as an additional control we find that the Pum NTD has no effect in the absence of the Nos NTD; both Pum RBD and PumC[889] recruit the isolated Nos RBD to an equivalent extent (“n.s.” in Fig. 2A).

**Figure 2.**
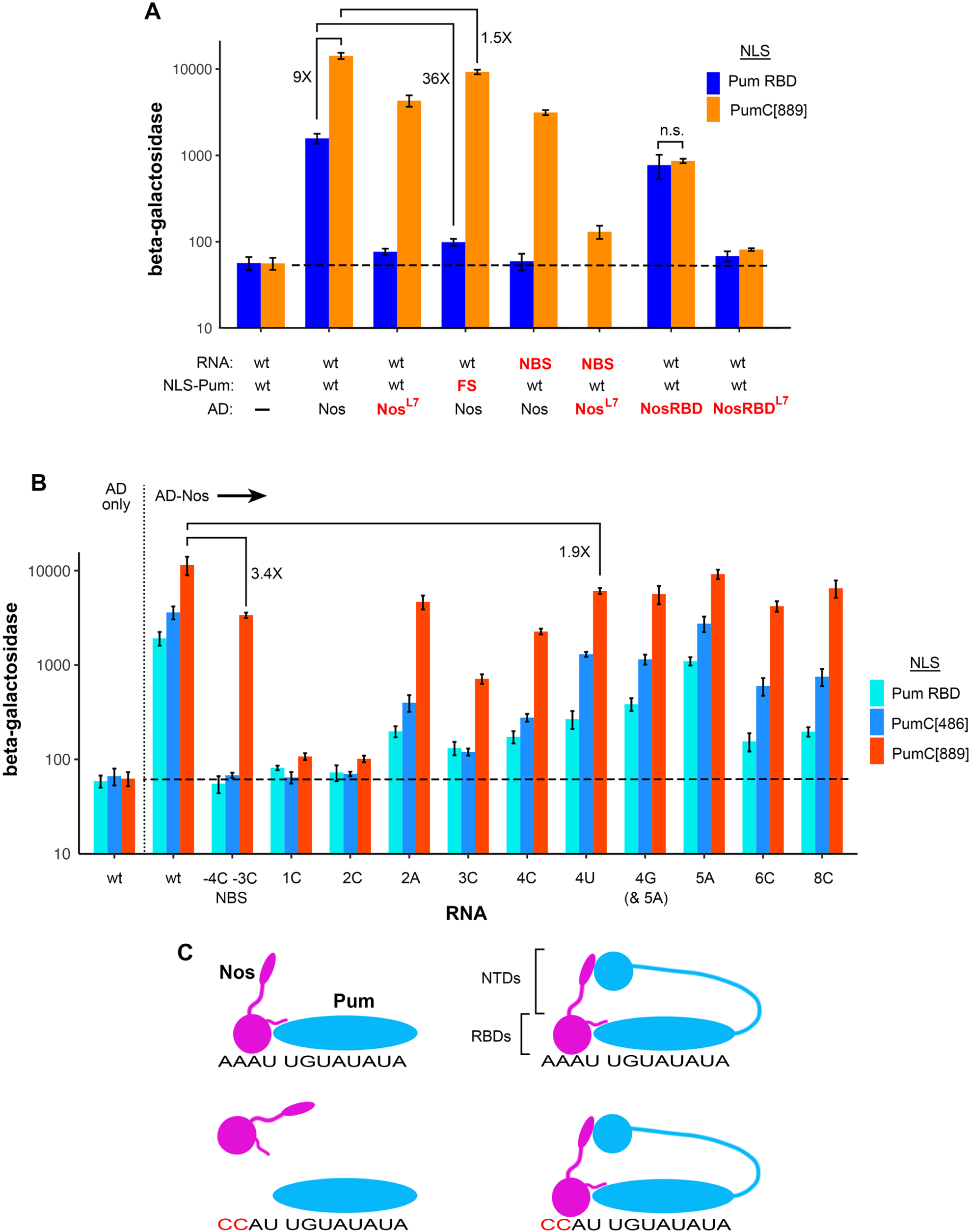
The Pum NTD alters Nos recruitment. A. Validation of the 4-hybrid assay used to test Nos recruitment. Logarithmic plot of the results of β-galactosidase assays of various 4-hybrid experiments, comparing the ability of NLS fusions of the Pum RBD and PumC[889] to recruit Nos to a NRE. The three vector-encoded components are indicated below the plot, with mutant derivatives in red. In the pair of controls at the left, yeast expressed AD only from an empty vector. The low basal activity in the absence of Nos further demonstrates that RNA-bound PumC[889] does not fortuitously activate transcription of the *lacZ* reporter. As noted in the text, the activity of PumC[889] better accounts for the activity of the NBS mutant in vivo; it is also consistent with the observation that activity of the Nos^L7^ mutant is reduced but not eliminated in vivo (13,25), although the extent of this reduction has not been quantified. Activity of the F1367S mutant has not been tested in vivo, to our knowledge. B. As in Fig. 1, logarithmic plot of the results of β-galactosidase assays measuring Nos recruitment by three NLS-fusions to Pum (color coding to the right) to the same NRE derivatives as in Figure 1 in yeast 4-hybrid experiments. The empty vector “AD only” control at the left shows the basal level of β-galactosidase. C. Model of the contribution of the Nos and Pum NTDs to recognition of the wild type NRE (above) and the NBS mutant (below). The NTDs are likely unstructured, and represented by extended chains and globules. In the absence of the Pum NTD (left), recruitment of Nos is dependent solely on the network of interactions between the Nos and Pum RBDs, as well as interactions between Nos and the RNA (i.e., as shown in Fig. 1A). In the presence of the Pum NTD (right), the elevation in apparent affinity and protein-protein interaction with the Nos NTD permit recruitment of Nos even to the NBS mutant, with only a 3.4-fold reduction in binding.

We next asked whether the presence of the Pum NTD alters the sequence-specificity of ternary complex formation by, in essence, repeating the experiment of Fig. 1 but measuring recruitment of full-length Nos rather than binding of Pum. To a first approximation, the NTD alters Nos recruitment in the same manner as it alters Pum binding, markedly relaxing specificity. Substitutions at positions +4, +6, and +8 (as well as the +2A substitution) are well tolerated; in fact, Nos is recruited to these mutant sites to a greater extent than to the NBS mutant, a useful benchmark for activity in vivo, as described below (Fig. 2B). Additionally, we note that PumC[486] recruits Nos more efficiently than the Pum RBD (but not as efficiently as PumC[889]), suggesting that residues C-terminal to the Pum RBD play a secondary role in Nos recruitment.

To our knowledge, no measurements of ternary complex formation by native Nos and Pum have been described. However, two observations suggest that the activity of full-length Pum in vivo is better modelled by the properties of PumC[889] than by the isolated RBD. First, as described above, both canonical UGUA (e.g., wt = 4A) and non-canonical UGUU (i.e., 4U) sites mediate Nos+Pum activity in ovaries and embryos (3), consistent with the efficient recruitment of Nos to the 4U “mutant” by PumC[889] (Fig. 2B). Second, the NBS mutant NRE retains substantial activity in the embryo, mediating repression of maternal *hb* mRNA by Nos and Pum such that 2-3 abdominal segments (of a full complement of 8) develop on average (14). This observation is consistent with the modest 3.4-fold reduction in Nos recruitment observed with PumC[889] but not the abolition of recruitment observed with the Pum RBD (Fig. 2B).

Taken together, the data in Figs. 1 and 2 are consistent with the model for Nos+Pum binding to the NRE shown schematically in Fig. 2C. In the case of the isolated RBDs, the cooperative network of weak protein-protein and Nos-NBS interactions (i.e., Fig. 1A) bring the RBDs together on the NRE, stabilizing a ternary complex (4,5). Deleterious substitutions that affect any component of the network, such as mutations in the NBS, eliminate ternary complex formation. We show below that the Nos and Pum NTDs interact without binding RNA (see Fig. 4), providing an independent mechanism to bring their respective RBDs together. As a result, substitutions in the NBS (for example) only modestly reduce ternary complex formation with Nos and PumC[889].

In the experiments described in Figs. 1 and 2, we studied binding to a model wild type NRE sequence that showed high affinity for both the RBD and Nos+RBD in preliminary experiments. The model site is, in essence, a chimera with the NBS of *hb* NRE2 juxtaposed to the PBS of NRE1 (Fig. 3A). We wished to test Pum binding and Nos recruitment to authentic NREs (i.e., sites shown to mediate regulation in vivo), particularly in view of conflicting reports in the literature. For example, the RBD has been shown to bind on its own to the well-characterized *CycB* NRE in one report but not another (5,19).

**Figure 3.**
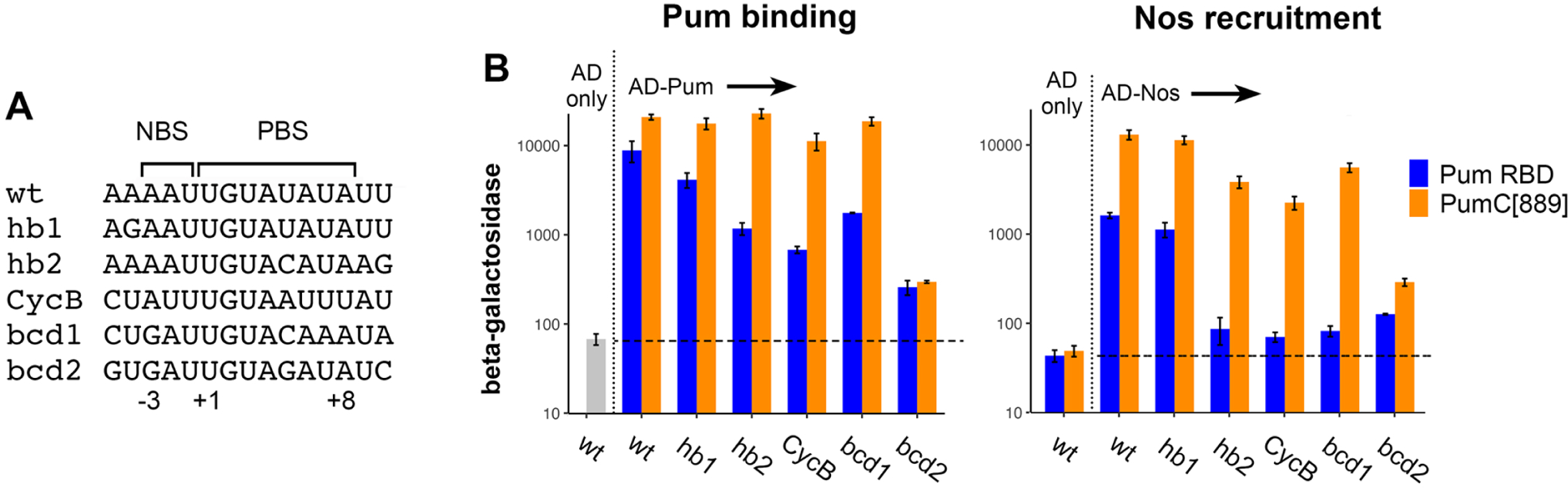
Joint binding of Pum and Nos, each bearing its NTD, correlates with the activity of authentic NREs in vivo. A. Sequences of the sites tested. B. As in Figures 1 and 2, logarithmic plots of the results of β-galactosidase assays measuring Pum binding and Nos recruitment by either the Pum RBD or PumC[889]. Empty vector controls at the left, as in previous figures. Note that the *bcd* NRE has been defined functionally as a 45 nt sequence bearing both of the Pum binding motifs shown here (13), and that the individual contribution of each to regulation in vivo has not been assessed in vivo, to our knowledge. The observation that Nos is recruited more efficiently to *hb1* NRE than *hb2* NRE is consistent with an earlier report showing that the *hb1* site confers greater repression in embryos (17).

Accordingly, we assayed RNA-binding and Nos-recruitment by either the isolated Pum RBD or PumC[889], comparing binding to the model wild type NRE with binding to: the two NREs in *hb*, the *CycB* NRE, and two fragments of the *bcd* NRE (Fig. 3A). As shown in Fig. 3B, relative to binding of PumC[889], binding of the Pum RBD is reduced ∼ 2-fold to the *hb1* NRE, but between 5- and 15-fold to the remaining authentic sites. (The *bcd2* sequence bears a non-consensus G residue at position 5 and has not been shown to play a role in regulation in vivo.) The NTD substantially stimulates interaction with all the authentic sites, such that PumC[889] binding to the *hb1*, *hb2* and *bcd1* NREs is indistinguishable from binding to the model NRE, and binding to the *CycB* NRE is reduced < 2-fold. With respect to Nos recruitment, the RBD recruits Nos to the *hb1* NRE, but barely recruits Nos above background to the other authentic sites. The NTD stimulates Nos recruitment between 10- and 68-fold to all the authentic sites, including the *CycB* NRE. We conclude that the Nos and Pum NTDs play a substantial role in joint binding to NREs that have been shown to regulate translation in vivo.

### Pum and Nos NTDs mediate hetero- and homotypic interactions

The results outlined above could be explained if interactions between the Pum and Nos NTDs, independent of interactions between the RBDs, stabilize the Nos/Pum/NRE ternary complex, as indicated schematically in Fig. 2C. To directly test this idea, we performed yeast 2-hybrid experiments, the results of which are shown in Fig 4.

**Figure 4.**
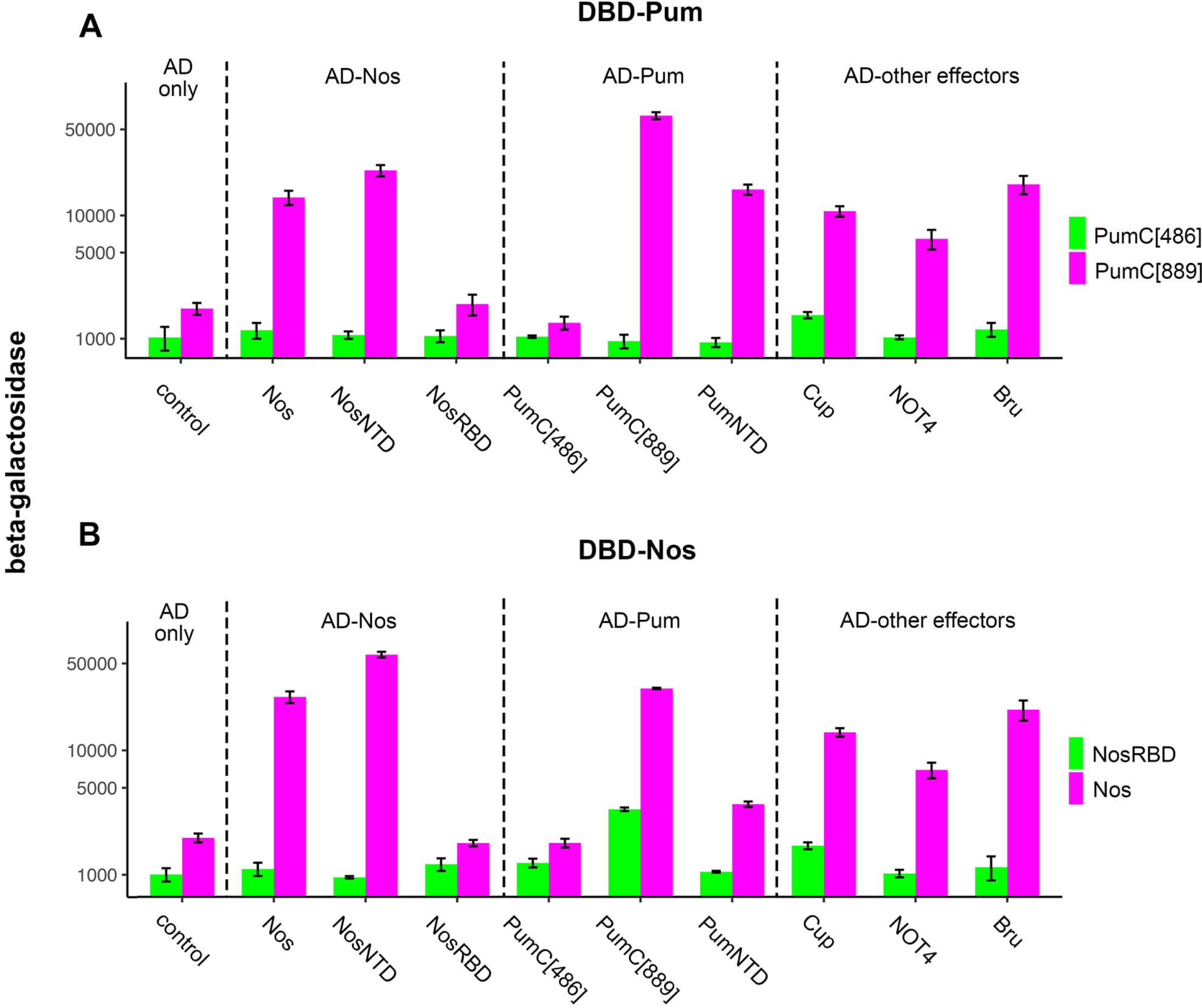
The Pum NTD interacts with itself and with the NTD of Nos. Logarithmic plots of the results of β-galactosidase assays of various 2-hybrid experiments, measuring interactions between various protein pairs as indicated. In (A), the DBD-fusion “bait” is either PumC[486] as a control lacking the Pum NTD or PumC[889], as indicated to the right. AD-fusion “prey” proteins are indicated above the plot. In (B), the DBD-fusion “bait” is either full length Nos or the Nos RBD, as indicated to the right. We were unable to test binding to a DBD-fusion with the isolated Nos NTD, since it auto-activates β-galactosidase expression in the absence of an AD-fusion partner. Note that the background level of β-galactosidase is higher in the yeast strain used in these experiments than in the strain used for 3- and 4-hybrid RNA-binding experiments elsewhere.

We first tested interaction of the Pum NTD with AD fusions to three sets of proteins: various derivatives of Nos, various derivatives of Pum, and various effectors thought to mediate regulation by Nos. In Fig. 4A, we expressed DNA-binding domain (DBD) “bait” fusions to PumC[889] and PumC[486], which bear and lack the NTD, respectively. As shown in Fig. 4A, PumC[889] interacts with full-length Nos and the isolated Nos NTD, but not with the Nos RBD. The NTD in PumC[889] also mediates homotypic interactions with the Pum NTD and with three proteins implicated in Nos-dependent regulation from previous work--Cup, NOT4, and Bruno (Kadyrova et al., 2007; Verrotti & Wharton, 2000; Marhabaie et al., 2023). PumC[486], which lacks an NTD, does not exhibit substantial interaction with any of the tested AD-fusion “prey” proteins.

We next tested the same set of protein-protein interaction, but with the DBD- and AD-fusion partners swapped. As shown in Fig. 4B, the Nos NTD mediates interaction with full length Nos and the isolated Nos NTD, but not with the Nos RBD. The Nos NTD also mediates heterotypic interactions with Pum and with the three known effector proteins (Fig. 4B). On its own, the Nos RBD exhibits negligible interactions with all factors except PumC[889], with which it interacts weakly (3.3-fold above background).

In summary, the 2-hybrid experiments described above support the model in Fig. 2C, in which interactions between the Pum and Nos NTDs off the RNA supplement the network of RNA-proximal interactions between RBDs to stabilize Nos/Pum/NRE complexes.

### NTDs allow binding to tandem mutant NREs

In all the experiments above, we measured binding to RNAs with a single NRE. We wondered whether the hetero- and homotypic interactions between Pum and Nos NTDs might enhance binding to an RNA with multiple suboptimal NREs, which would further expand the repertoire of mRNAs targeted by Nos+Pum.

To test this idea, we returned to yeast 4-hybrid experiments to measure recruitment of various AD-Nos fusions to RNAs with tandem NREs (Fig. 5). The key result is highlighted in blue boxes below the bargraph: when both Nos and Pum proteins possess their respective NTDs, Nos is recruited to essentially the same extent to (1) an RNA with a single wild type NRE as to (2) an RNA with <u>no</u> wildtype NREs, in which tandem mutant NREs supply non-adjacent binding sites for Nos and Pum. Nos recruitment to this RNA with tandem mutant NREs is relatively efficient, with β-galactosidase levels 35% of those for recruitment to the control RNA with tandem wildtype NREs. Furthermore, Nos binding to the tandem mutant NREs is dependent on both the Pum and Nos NTDs, the C-terminal Nos tail that mediates interaction between the Nos and Pum RBDs, and the presence of a wildtype NBS and PBS on the RNA.

**Figure 5.**
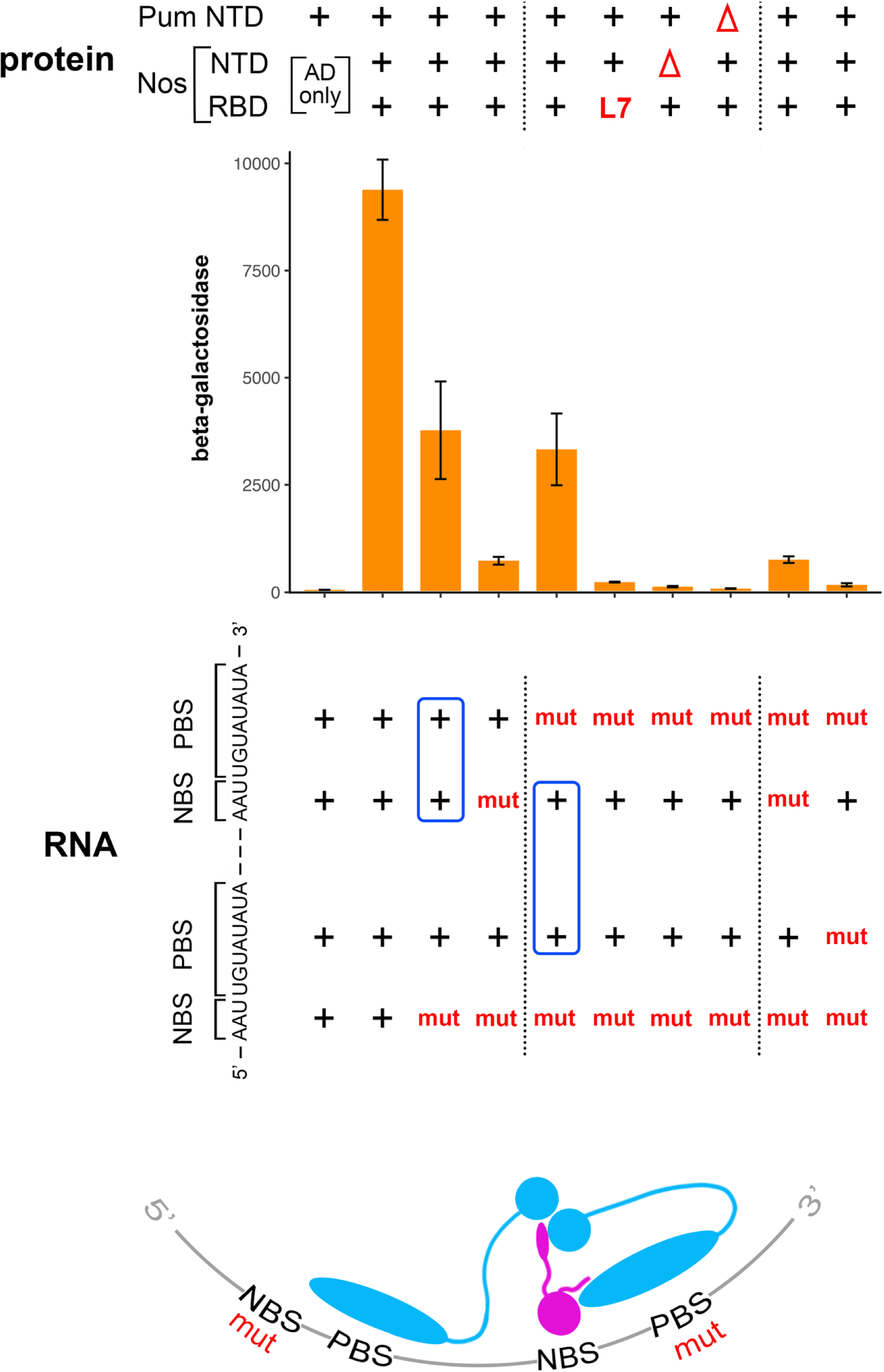
The Pum and Nos NTDs allow recruitment of Nos to an RNA with two suboptimal, mutant NREs. At the top the components in each 4-hybrid experiment are shown. The Pum RBD is wt in each case, and the first line indicates whether its associated NTD is present (+) or absent (Λ1) (i.e., PumC[889] vs. PumC[486]). The second and third lines indicate the identity of the Nos moeity in each case, in particular, whether its NTD is present (+) or absent (Λ1) (i.e., full-length Nos or Nos RBD only), and whether its RBD is wild type (+) or bears the L7 mutant. The bargraph is a linear plot of β-galactosidase assays measuring Nos recruitment to RNAs with various tandem NREs. The linear plot was chosen here to emphasize the difference between the RNAs highlighted in blue boxes, which is obscured in a logarithmic plot. Note that, when displayed on a linear plot, the error bars appear larger in this experiment than in others in this report (which are displayed on logarithmic plots). The sequence of the NREs in the fully wild type RNA is below the bargraph to the left, followed by a schematic indication of whether the NBS or PBS element is wild type (+) or mutant in each NRE variant. The critical comparison between an RNA with one intact NRE (lane 3) and an RNA with no intact NREs but separated wild type NBS and PBS elements (lane 5) is highlighted in blue boxes. At the bottom is a schematic interpretation of binding in the experiment of lane 5. Pum bound upstream recruits Nos at a distance via cooperative interactions among NTDs that allow assembly of a Nos+Pum complex downstream, even though the downstream PBS bears a deleterious 3C mutation indicated by “mut.” In the control in lane 10, the upstream PBS bears a deleterious 1C mutation; and in lanes 5-10, the downstream PBS bears a deleterious 3C mutation. Where indicated, the mutant NBS bears the −4C & −3C substitutions assayed in Figures 1 and 2.

Our interpretation of the experiments is shown at the bottom of Fig 5. In essence Pum bound upstream can drive formation of a Nos/Pum/RNA complex “at a distance” downstream, even when the downstream 3C mutant site has only a low affinity for Pum (e.g., as in Fig. 1). Although we favor this model, we cannot rule out an alternative in which a novel deployment of the Nos C-terminal tail, mediates interaction between Pum bound to the upstream PBS and Nos bound to the downstream NBS.

We note that a bioinformatic analysis of the key RNA sequence in these experiments would not identify it as a likely target for Nos+Pum, since it bears no intact NREs. The results of Fig. 5 suggest that the NTDs of Nos and Pum may expand the repertoire of mRNA targets beyond our current understanding, although we have not tested whether such sequences can mediate regulation in vivo.

### The human Pum2 NTD also alters RNA-binding

We next asked whether the NTD of human Pum2 has a similar effect on RNA-binding. The DmPum and HsPum2 RBDs are 80% identical overall, have identical RNA base-contacting residues, and bind to the same PBS consensus site in vitro (27). In contrast, the NTDs of the two proteins both consist of low complexity sequence predicted to form intrinsically disordered regions (IDRs), with four amino acids-- Ala, Gln, Gly, and Ser-- making up ∼55% of both domains. Excluding low-complexity regions, the two NTDs share < 10% sequence identity.

We expressed either the HsPum2 RBD or a longer fragment bearing 384 residues of its NTD fused to the RBD with the panel of NREs shown in Fig. 1, and measured RNA-binding in 3-hybrid experiments as described above. As shown in Fig. 6, the human and Drosophila NTDs appear to alter activity of their respective RBDs in a similar manner. Binding specificity outside the UGU residues at +1 to +3 is significantly relaxed, with binding of Hs Pum2 bearing its NTD to the 4U, 4G, 6C and 8C sites reduced at most twofold relative to binding to the wild type site (Fig. 6). In contrast, binding of the RBD to the 4U, 4G, 6C and 8C sites is reduced from 4.5- to 21-fold. Finally, binding to the wild type site is enhanced 1.8X by the NTD, even though the NTD-bearing protein is expressed at a lower level than the isolated RBD (S6 Data). Thus, the NTDs of human and fly Pum have similar effects on the activity of their respective RBDs, despite limited homology (see S3A Fig).

**Figure 6.**
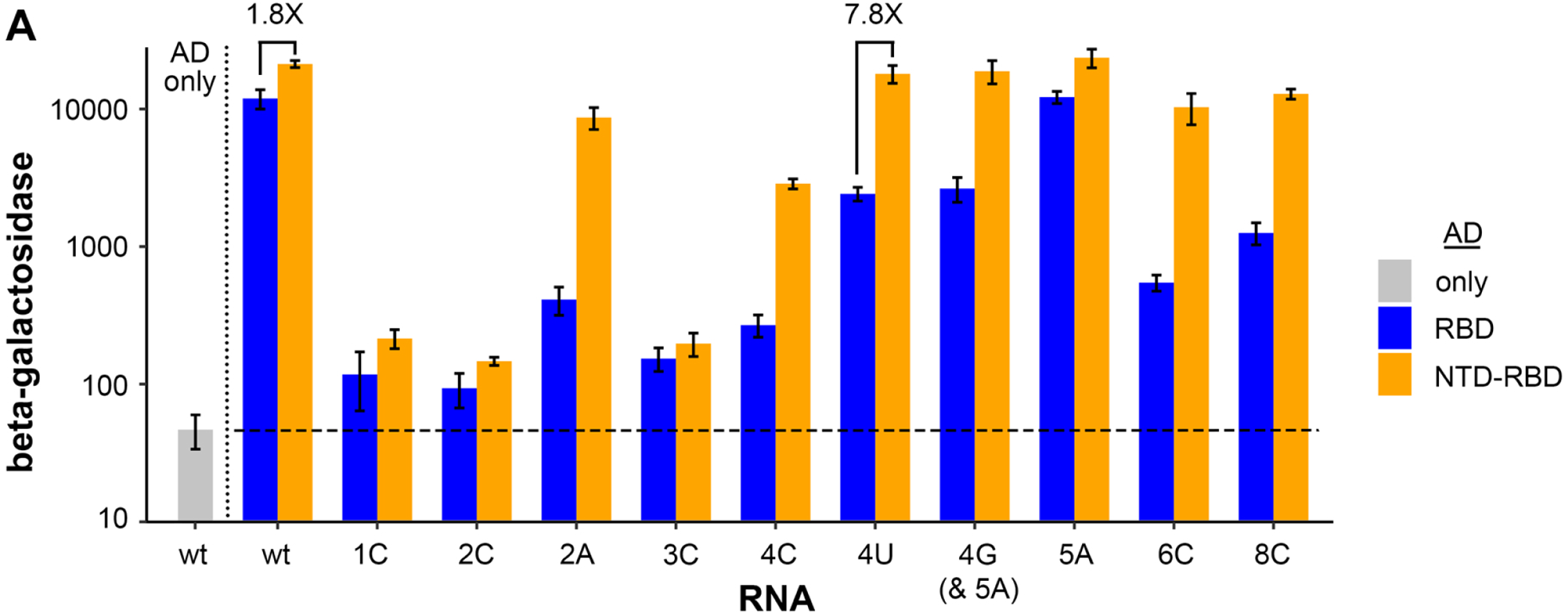
The human Pum2 NTD alters binding specificity of its RBD. Logarithmic plot of β-galactosidase 3-hybrid assays measuring binding to the *hb* NREs assayed in Figs. 1 and 2 (with the exception of the NBS mutant) to either the HsPum2 RBD or a fragment of HsPum2 bearing 384 residues of its NTD fused to the RBD, as indicated. At the left is a negative control from yeast expressing AD only.

### NTD-dependent alteration of Pum binding in vivo

The NTD in PumC[889] consists largely of low-complexity sequence thought to comprise an intrinsically disordered domain. We wondered whether the ability to alter RNA binding maps to a discrete sub-region of this NTD. As an initial step in addressing this question, we prepared two additional Pum derivatives, PumC[889β] and PumC[684], which bear the amino- and carboxy-proximal halves of the NTD, respectively, fused to the RBD (S3A Fig). We then tested binding to various NREs in yeast 3-hybrid experiments.

We find that PumC[889] and PumC[889Δ] have similar relaxed specificities, recognizing the 4U, 4G, 6C, and 8C mutant NREs with only between 1- and 2.5-fold differences in binding (Fig. 7 S1). In contrast, PumC[684] distinguishes wild type from mutant bases at positions +4, +6, and +8 of the NRE with slightly greater selectivity than does the isolated RBD (S3B Fig). For example, PumC[684] and the RBD bind at 31- and 12-fold lower levels, respectively, to the 4U site than to the wt NRE. Thus, residues that relax binding specificity map to the 206 residue, amino-proximal portion of the NTD in PumC[889]. Half of this region consists of sequence rich in Ala and Gln residues (56%), and the other half bears a short motif present in both the Hs Pum1 and Hs Pum2 NTDs (S3A Fig).

**Figure 7.**
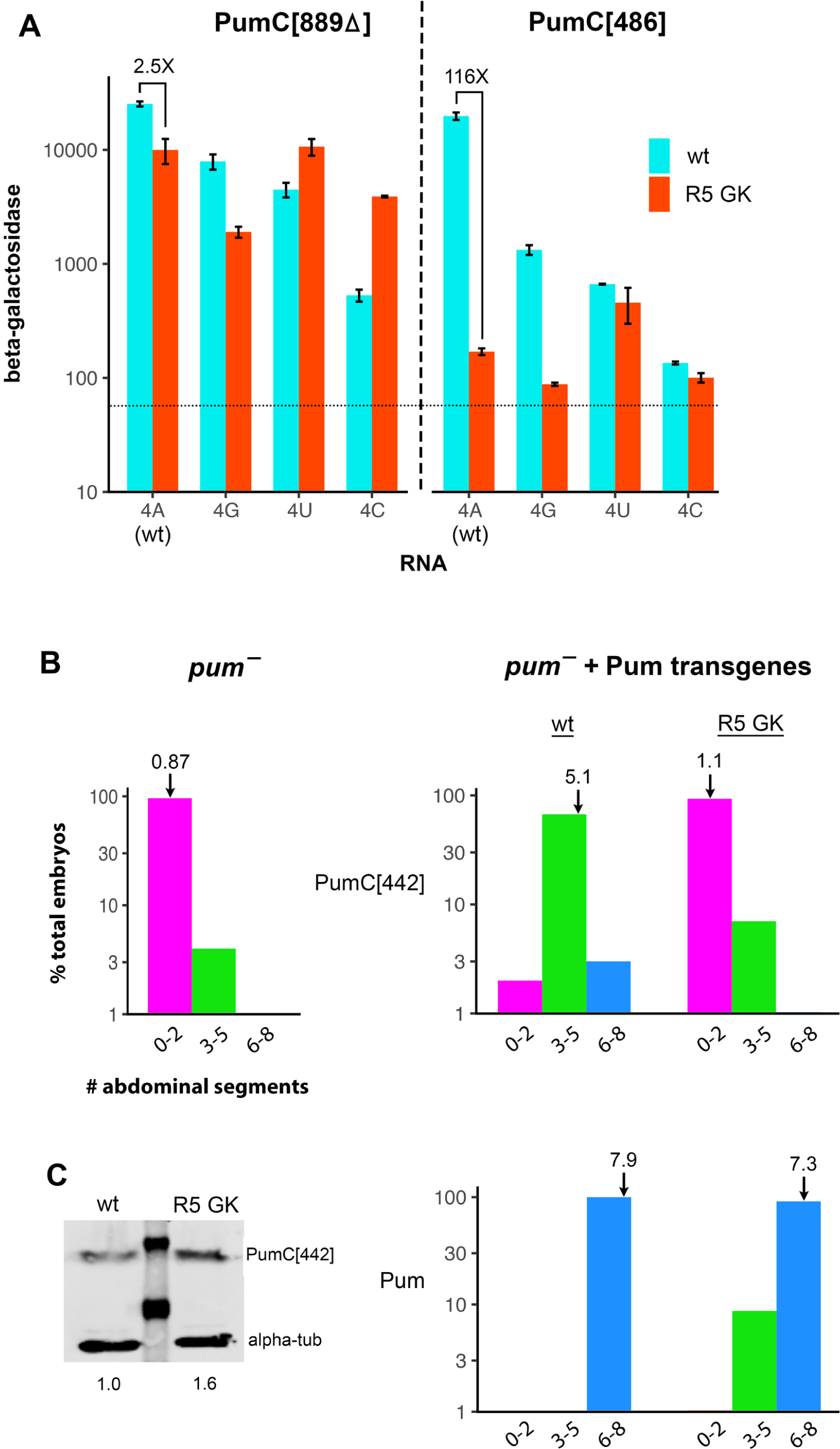
The Pum NTD is essential for activity of a variant RBD in yeast and in embryos. A. Logarithmic plot of yeast 3-hybrid experiments comparing the binding of Pum derivatives with (PumC[889Δ]) and without (PumC[486]) the critical region of the NTD that alters binding specificity. As indicated above and to the right, the RBD is either wild type or bears Gly and Lys substitutions in the fifth Puf repeat (R5 GK), which recognizes position +4 in the NRE. Binding is assayed to NREs with each of the four bases at position +4, as indicated on the x-axis. In these experiments, each RNA bears an A rather than a U at position +5 to avoid a potential cryptic Pum binding site in the 4G mutant. The horizontal dashed line is the mean AD-only control from similar experiments elsewhere in this report. B. Pum-dependent regulation of maternal *hb* mRNA governs abdominal segmentation. Abdominal segmentation was scored among embryos from *pum* hypomorphic mutant females (left) or *pum* mutant females that also carry one of four Pum transgenes, as indicated (panels on right). Transgenes on the top row encode PumC[442], which bears residues 1093-1533 of Pum, containing the RBD and C-terminal residues. Transgenes on the bottom row encode full-length Pum. Embryos from females bearing null alleles of *nos* or *pum* develop 0 abdominal segments, whereas embryos from wild type females develop 8 abdominal segments. The x-axis in each plot displays the number of abdominal segments (binned) and the (logarithmic) y-axis displays the % of total embryos in each bin. In each bargraph, the average number of abdominal segments is indicated with an arrow. C. Western blot showing maternal expression of PumC[442] in 0-2 hr embryos, with a lane containing MW markers separating wild type (left) and R5 GK mutant (right) proteins. The relative amount of each Pum protein is shown below. Transgenic Pum proteins were Myc-epitope tagged near the C-terminus and detected with anti-Myc antibody, with endogenous α-tubulin as a loading control. The visible MW markers are 98 and 64 kD.

In work to be described elsewhere, we have isolated mutant derivatives of PumC[889Δ] that use non-wild type amino acids to recognize various NREs. Here we use one of these mutants to test the role of the NTD in RNA binding in vivo, as explained below.

The fifth Puf repeat (R5) of the wild type RBD bears Cys and Gln residues that make sequence-specific contacts to the fourth position of the PBS (4A in the wild type NRE). We initially isolated the R5 GK derivative (i.e., bearing Gly and Lys base-contacting residues) of PumC[889Δ] in a screen for mutants that bind a 4U site. Subsequent analysis revealed that the R5 GK protein also recognizes the wild type 4A NRE efficiently, with binding reduced only 2.5-fold in comparison to the wild type protein (Fig. 7A). In addition, R5 GK exhibits significantly relaxed specificity, binding 2.4- and 7.3-fold <u>better</u> to the 4U and 4C mutant sites, respectively, than wild type PumC[889Δ].

We also tested binding of an R5 GK derivative of PumC[486], which led to a remarkable observation: binding of R5 GK to the wild type (4A) NRE is essentially dependent on the NTD. As shown in Fig. 7A, in the absence of the NTD, binding of R5 GK to the 4A site is reduced 116-fold in comparison to the wild type PumC[486] protein. Based on this finding, we realized we could use the R5 GK mutant to test whether the NTD alters Pum binding in the Drosophila embryo as follows.

During early embryogenesis, joint regulation of a single mRNA by Nos and Pum-- maternal *hb* -- is necessary and sufficient to allow development of abdominal segmentation (28–30). This regulation is mediated by two NREs in the *hb* 3’-UTR, each of which bears a canonical A at position +4 (sequences shown in Fig. 3A). In embryos from females bearing a *pum* hypomorph, maternal *hb* is not repressed and the embryos develop on average < 1 abdominal segment (Fig. 7B). In this genetic background, previous work has shown that abdominal segmentation is partially rescued by maternal expression of a C-terminal fragment of the protein that lacks the NTD (PumC[442]) and completely rescued by expression of full-length wild type Pum (14). As shown in Fig. 7B, we find that wild type PumC[442] rescues development of 5.1 abdominal segments on average (of the full complement of 8), and full-length Pum rescues development of 7.9 abdominal segments.

The properties of the R5 GK mutant in yeast binding experiments make a clear prediction: it should rescue the abdominal segmentation defects of *pum*^−^ embryos, but only with the Pum NTD in cis. As shown in Fig. 7B, this is what we observe. In the absence of the NTD, the R5 GK mutant does not significantly rescue abdominal segmentation above the low level in the *pum* mutant background (*p* = 0.18 by Fisher’s Exact Test). In contrast, with the NTD in cis (as part of the full-length protein), R5 GK drives almost complete abdominal segmentation (7.3 segments on average), only slightly less than the 7.9 segments (on average) in embryos rescued by Pum with the wild type RBD. The R5 GK mutant derivative of PumC[442] is expressed to a slightly higher level (1.6-fold) than wild type PumC[442] in early embryos (Fig. 7C), and thus its inactivity is not due to reduced expression or stability. Finally, we note that although the full-length GK mutant efficiently rescues abdominal segmentation, most of the resulting larvae are inviable, with few surviving to third instar (< 1%). In contrast, the majority (> 50%) of *pum*^−^ embryos rescued by the wild type full-length control protein survive to third instar or beyond. A likely explanation is that the GK mutant has relaxed specificity (as shown in the yeast experiments of Fig. 7A) and therefore inappropriately regulates a number of mRNAs in embryos, especially those bearing NRE-like sequences with a C at position +4.

In conclusion, the NTD likely promotes binding of the R5 GK mutant to the *hb* NREs in the embryo via both of the activities identified in this report-- (1) altering recognition of the RNA by the RBD in cis, and (2) mediating protein-protein interactions that stabilize Nos/Pum complexes on the NREs.

## Discussion

Previous work on RNA recognition by Pum and Nos has focused on studies of their RBDs. Here we show that the NTDs of both proteins play important roles in RNA binding and propose models for how they act.

The Pum and Nos NTDs interact with each other, which enhances joint occupancy of the wild type NRE by Nos and Pum 6- to 9-fold (Fig. 2). We suggest that weak Nos/Pum hetero-oligomerization mediated by their NTDs promotes binding to specific NREs by a conventional mechanism-- increasing the avidity for RNA via protein-protein interactions that elevate the probability of joint occupancy by both RBDs. A similar strategy is employed by members of the RRM-domain family, many of which have multiple low-affinity RRMs. For example, as an isolated domain, the first RRM of Xenopus PABP does not bind detectably to poly(A) and the second binds with only modest affinity (31). However, a fragment of the protein bearing RRM1+RRM2 binds with high avidity, as does a similar fragment of the closely related human PABP (32).

A second, less conventional role for the Pum NTD is revealed by two sets of experiments in which it alters the binding specificity of the RBD in cis. First, the NTD relaxes binding specificity of the wild type protein at positions +4, +6, and +8 of the NRE (Fig. 1); the most extreme example is the 25-fold elevation of binding to the 4U mutant site conferred by the NTD. Second, the NTD alters base discrimination at position +4 for the R5 GK mutant RBD, enhancing binding to the wild type 4A site ∼ 60-fold. How might the NTD alter RNA recognition?

Structural and biochemical studies provide precedent for modification of Puf domain activity by other proteins in trans; we suggest that the Pum NTD performs a similar function, but does so by acting in cis. One case of Puf activity modification is effected by the Nos RBD, which relaxes the specificity of Pum at NRE positions +5 through +8 (5). Interactions of Nos with positions −3 to −1 of the NRE and with the C-terminal end of the Pum RBD cause local conformational changes in the protein. Pum remains in contact with 3’-proximal bases in the NRE, and it is currently unclear how specificity at these positions is relaxed. A second case is modification of C. elegans FBF-2 binding by LST-1, which binds near the carboxy-terminus of the FBF-2 Puf domain, on its “outside,” non-RNA contacting surface (33). Like Nos, LST-1 relaxes binding specificity for bases at the 3’-end of the binding site, which remain in contact with the Puf domain. The Pum NTD could interact with its RBD, the RNA, or both to alter binding specificity. Relatively weak interactions might suffice, since the interaction would be intramolecular.

We note that both the inter- and intra-molecular activities of the Pum and Nos NTDs may expand the repertoire of regulated mRNAs beyond those bearing canonical NBS+PBS sequences. As described above, relaxation of binding specificity mediated in cis by the Pum NTD appears to account for the efficient targeting of 4U “mutant” NREs by Nos and Pum in vivo (3). Interactions in trans between the Nos and Pum NTDs are evidently stronger than those between the RBDs; the former are easily detected in yeast 2-hybrid experiments (Fig. 4), while the latter are observed only when the RBDs are aligned and brought together on the NRE in a ternary complex (Fig. 2). We therefore suggest that the NTDs likely play a major role in site selection in vivo. In Fig. 5, we show that Pum can recruit Nos relatively efficiently to an RNA with no intact NRE, as long as the NTD of both proteins is intact. It will be important to test whether such RNA sequences can mediate regulation in vivo with endogenous proteins.

To further test the model for NTD activity outlined above, we would like to measure binding in vitro with purified proteins. However, to date we have been unable to express soluble versions of either (1) Pum bearing both the RBD and the critical portion of the NTD or (2) Nos with its NTD. Perhaps further definition of the residues that mediate interaction will permit preparation of smaller, soluble derivatives, although the properties of IDRs present a notorious challenge. A further consideration for future studies is that amino acid residues 1-643 of Pum might affect the activities of the NTD that we focus on in this report (residues 644-1092).

In addition to its effects on RNA-binding, the Pum NTD may act as a scaffold for protein-protein interactions that regulate mRNA translation and stability, based on the results with Bru, Cup, and NOT4 shown in Fig. 4. The structure prediction program alpha-fold (34,35) identifies only one feature in the Pum NTD: a 35 residue alpha-helix consisting of amino acids 711-745 that is predicted with low- to moderate confidence. This observation is consistent with the idea that the NTD is largely disordered. It may therefore promote interaction with itself, with similarly disordered regions in the NTD of Nos, and with various translational regulators (e.g., Cup, NOT4, and Bru) via via liquid-liquid phase separation, rather than via distinct binding motifs for each partner. It is possible that Nos+Pum-bound RNA also contributes to aggregation, as has been suggested previously for RNA-binding proteins bearing aggregation-prone domains (36). Two other reports are consistent with these ideas. First, the Dm Pum NTD has been shown to aggregate when expressed in yeast (37). And second, when expressed in HeLa cells, the Hs Pum2 NTD drives the protein into stress granules, which are phase-separated RNA-protein condensates (38).

The human and fly Nos and Pum proteins share a number of key structural and functional features. The Zn fingers of the Nos proteins have similar sequences that underlie likely structural homology (39). The Puf domains are structurally similar and bind the same consensus RNA sequence 10/22/23 8:51:00 AM. Finally, data presented in this report show that the Dm Pum and Hs Pum2 NTDs modify the binding activities of their respective RBDs in a similar manner.

Despite these similarities, it is not clear whether the NTDs of the human proteins function as do the corresponding NTDs of the Drosophila proteins. Indeed, it is not yet clear whether the human Nos and Pum proteins bind together to regulate target mRNAs, as, to our knowledge, there is neither compelling genetic nor biochemical evidence for joint activity. The Zn finger domains of human Nos2 and Nos3 have been reported to bind jointly with human Pum1 and Pum2 to RNA in vitro, but at Nos concentrations of ∼ 10 µM where non-specific interactions with either protein or RNA (or both) are difficult to exclude (40). Regulation of target mRNAs by murine Nos2 has been shown to be mediated by the RRM protein DND1 in the male germline (41). Thus, there exists at least one Pum-independent mechanism to recruit human Nos2 to RNA. Further work will be required to determine whether the NTDs of the various human Nos and Pum proteins function in vivo in an analogous manner to the corresponding domains of Drosophila Nos and Pum.

## Materials and Methods

### Drosophila strains and methods

Transgenic lines were constructed by microinjection of *w*^1118^ embryos using standard methods. The *pum*^−^ flies for the experiments in Fig. 7 were trans-heterozygotes of *pum*^Msc^ and *pum*^ET3^. We observed only minor differences among three or more transgenic lines, which were tested in preliminary experiments for rescue of abdominal segmentation. Fly stocks were maintained by standard methods and grown at 25°C. Embryos were collected on apple-juice agar plates smeared with yeast paste. Embryonic cuticle was examined by harvesting embryos 24 hours after removing adults, dechorionating with bleach, and mounting in Hoyers/lactic acid. Slides were cleared by heating for several hours and then examined by dark field microscopy using a Zeiss Axiophot. For the Western blot of maternal proteins in Fig. 7, 0-2 hr embryo collections were homogenized in sample buffer, electrophoresed on a SDS denaturing gel, and transferred to 0.2 µm nitrocellulose membranes by standard methods. The epitope-tagged Pum and α-tubulin proteins were detected by successive incubation with anti-Myc 9E10 monoclonal primary (SC-40, Santa Cruz Biotechnology) and anti-α-tubulin B-5-1-2 monoclonal primary (T5168, Sigma), followed by goat-anti mouse IRDye 680 goat anti-mouse IgG secondary antibodies (926-68070, LiCor), and detected on a LI-COR Odyssey CLx.

### Plasmids

Plasmids used in this work, and a description of their construction are listed in S8 Data. Note that the PumC[442] protein expressed in flies and described here is identical to Δ2’ in (14) with the exception of the 6X-Myc epitope tag near the C-terminus. The PumC[442] ORF begins at the native Pum AUG initiation codon and continues through the first 27 residues of the PA isoform, which are fused to the C-terminal 442 codons encoding the RBD and C-terminal residues. Note that the Nos RBD ORF described here is identical to Nos ΔN1 in (4) and bears, in addition to the Nos RBD the N-terminal 42 residues of Nos as well, which are required for efficient expression of the RBD in yeast for unknown reasons. For expression of RNA in 3- and 4-hybrid experiments, sequences inserted into pIII/MS2-2 were analyzed via m-fold (42) to ensure accessibility of each binding site.

### Yeast experiments

Measurements of β-galactosidase were performed essentially as described (3), from which the following description is taken. Yeast were transformed with the 2µ plasmids described in S8 Data by a standard lithium acetate/PEG protocol. For 2-hybrid experiments, we used the PJ69-4A strain (43), which is *MAT**a***, *trp1-901*, *leu2-3*, *112*, *ura3-52*, *his3-200*, *gal4*Δ, *gal80*Δ, *LYS2::GAL1-HIS3*, *GAL2-ADE2*, *met2::GAL7-lacZ*. Double transformants were obtained and grown on minimal SD dropout medium lacking tryptophan and leucine. For 3- and 4-hybrid experiments, we used the YBZ1 strain (44), which is *MAT**a**, ura3-52, leu2-3, 112, his3-200, trp1-1, ade2, LYS2::(LexAop)-HIS3, ura3::(lexA-op)-lacZ, LexA-MS2 coat (N55K)*. For 3-hybrid experiments, double transformants were grown on minimal SD dropout medium lacking uracil and leucine, and for 4-hybrid experiments triple transformants were grown on SD dropout medium lacking uracil, leucine, and tryptophan.

β-galactosidase activity was measured using β-Glo (Promega) essentially as described by Hook et al., 2005. Briefly, transformants were grown by diluting saturated overnight cultures into appropriate selective media to early-log phase (OD_600_ ∼ 0.3), and then 40 µl of culture were incubated with the same volume of β-Glo reagent for 60 minutes at room temperature in 96-well microplates. Four transformants were grown separately and assayed for each experiment, occasionally omitting a culture that grew unusually slowly or quickly. Samples were analyzed in a Veritas luminometer (Turner Biosystems Inc). The output signal from the luminometer was divided by the OD_600_ to normalize for the number of cells in each sample, and by 1000 (by convention) to generate a reading of β-galactosidase activity in arbitrary light units. *p*-values reported in Supporting Data are from unpaired t-tests.

Western blot samples were prepared by growing yeast cultures to OD_600_ = 0.3. Cells were harvested by centrifugation and resuspended in 50 mM Tris pH 7.4, 100 mM KCl, 0.1 mM EDTA, 0.1% NP40, and 1:50 protease inhibitor cocktail for yeast extracts (P8215, Sigma-Aldrich); lysed with the addition of glass beads (G-8772, Sigma) and 5 X 1 minute disruption in Mini-Beadbeater-8 (Biospec Products) at 4°; centrifuged at 15K for 10 min; and addition of SDS sample buffer to the supernatant. Western blots were performed as described above using monoclonal C29F4 rabbit anti-HA (3274, Cell Signaling Technology) to detect the HA-epitope tagged proteins encoded by plasmids and mouse monoclonal Sc Rpl3 (Developmental Studies Hybridoma Bank) to detect RpL3 (as a loading control). The anti-mouse secondary antibody from LiCor is described above, and the anti-rabbit IRDye 800 goat anti-rabbit IgG secondary antibody was 926-32211 from LiCor. *p*-values reported in Supporting Data are from unpaired t-tests.

## Supporting information

Wharton Supplemental Data 1

Wharton Supplemental Data 2

Wharton Supplemental Data 3

Wharton Supplemental Data 4

Wharton Supplemental Data 5

Wharton Supplemental Data 6

Wharton Supplemental Data 7

Wharton Supplemental Data 8

## Acknowledgements

We thank Drs. R Arvola, P Herman, and G Singh for comments on the manuscript, M Campbell for teaching RPW the rudiments of programming in R, the C Croce lab for sharing equipment, and M Wickens for sending yeast strain YBZ1. We acknowledge services of the OSUCCC Genomics Shared Resource which is supported by P30CA016058. The anti-RpL3 mouse monoclonal developed by John Warner was obtained from the Developmental Studies Hybridoma Bank, created by the NICHD of the NIH and maintained at The University of Iowa, Department of Biology, Iowa City, IA 52242.This research was supported in part by NIGMS R01GM084376 (RPW).

## Supporting Information

**S1 Fig.**
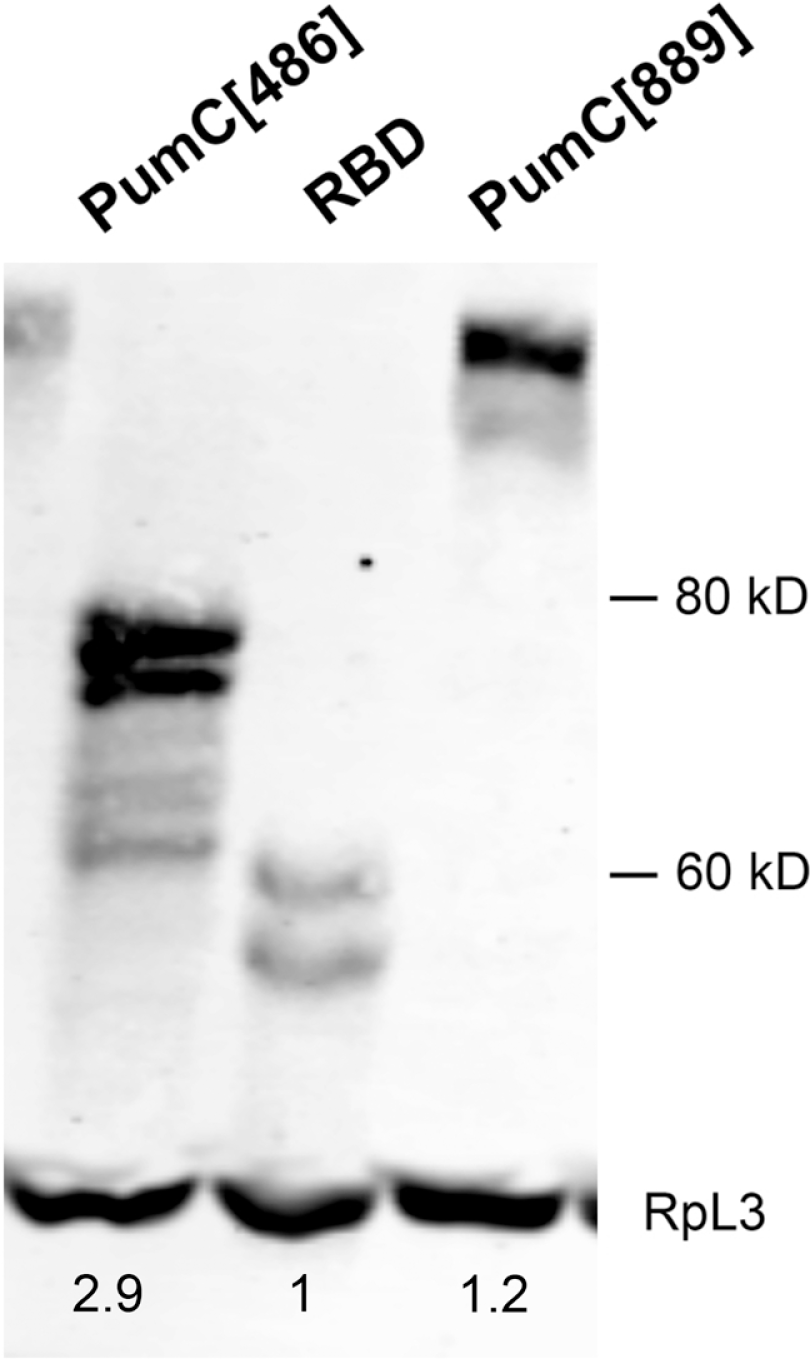
Western blot of AD-Pum fusions expressed in yeast. Each Pum protein is detected with a common, vector-encoded HA epitope tag. The ribosomal protein RpL3 is a loading control. We do not know whether the observed degradation occurs during growth of the culture or during lysis. The relative levels of Pum derivatives shown below are based on measurement of total HA-tagged protein. We also measured the relative amounts of the longest, presumptive full-length proteins (see S1 Data).

**S2 Fig.**
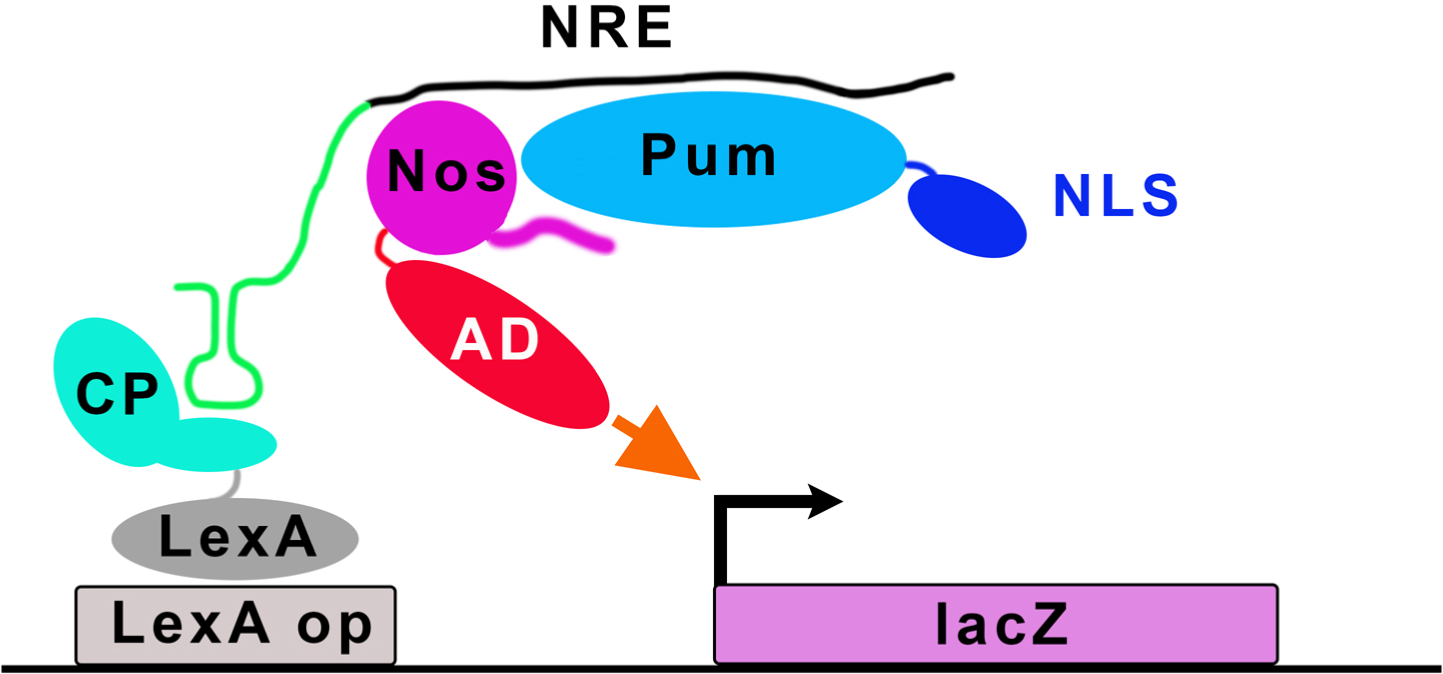
Schematic showing components of yeast 4-hybrid experiments that measure recruitment of Nos into a ternary complex with Pum and the NRE.

**S3 Fig.**
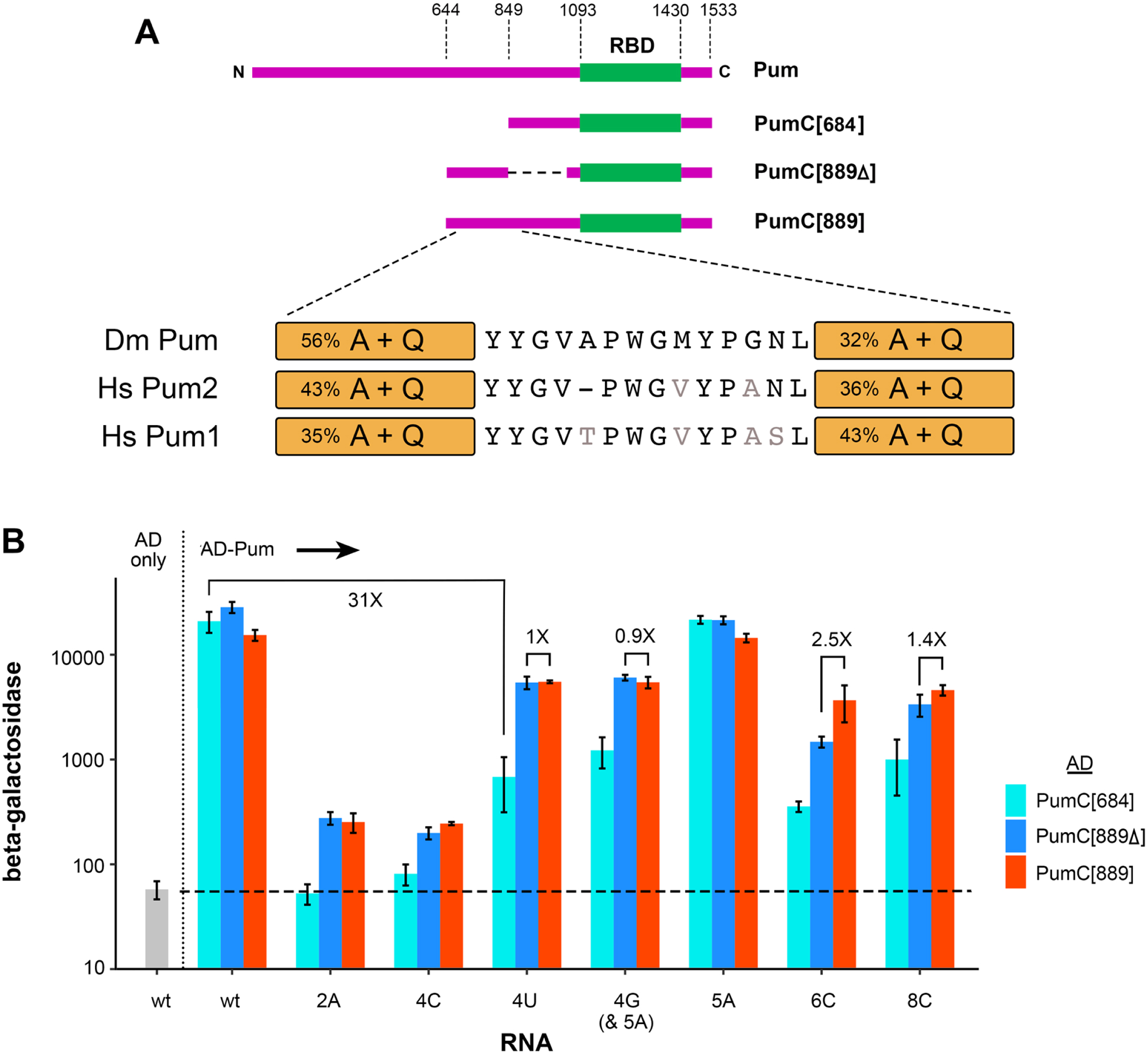
Relaxation of RNA-binding specificity maps to a sub-region of the Pum NTD. A. Schematic drawing to scale of the Pum derivatives tested below in B. The amino acid sequences below are the short motifs conserved among the Drosophila and human Pum NTDs, flanked by rectangles in which the Alanine and Glutamine content in the 100 adjacent residues is indicated. Similar but not identical residues in grey font. B. The region responsible for altering binding specificity maps to the proximal half of the NTD. Three-hybrid RNA-binding experiments to wild type (wt) and various mutant NREs of the three AD-Pum fusions indicated to the right.

**S1 Data. β-galactosidase measurements and statistical tests for Fig 1 and for AD-deleted derivatives of Pum fusions used in this report; quantitation and statistical tests for Western blots reported in S1 Fig.**

**S2 Data. β-galactosidase measurements and statistical tests for Fig 2 and for 3-hybrid binding of F1367 Pum derivatives to the wt NRE.**

**S3 Data. β-galactosidase measurements and statistical tests for Fig 3.**

**S4 Data. β-galactosidase measurements and statistical tests for Fig 4.**

**S5 Data. β-galactosidase measurements and statistical tests for Fig 5.**

**S6 Data. β-galactosidase measurements and statistical tests for Fig 6; quantitation and statistical tests for Western blot of AD-HsPum2 proteins.**

**S7 Data. β-galactosidase measurements and statistical tests for Figs 7A and S3; abdominal segmentation data for Fig 7B; quantitation and statistical tests for Western blots reported in Fig 7C**.

**S8 Data. Plasmids used in this work.**

